# Metataxonomic insights into the effects of enhanced nitrogen addition on the bacterial communities in the tropical paddy soil of West Bengal, India

**DOI:** 10.64898/2025.12.02.691912

**Authors:** Shuchishloka Chakraborty, Pinaki Sar

## Abstract

In relation to global rice cultivation, nitrogen (N) is fundamental for paddy soil and crop productivity; however, the exploitation of synthetic N fertilisation in the global agri-food system has emerged as a major environmental concern. In this study, the effects of various synthetic N-compounds on paddy soil bacterial communities, including species abundance, diversity, and interactions, were investigated under four different N-amended [NO2-, NO3-, NH4+, and urea] conditions using a microcosm-based culturomics approach. The N-amendments altered soil properties, adversely impacting microbial abundance, diversity, and community composition. The 16S rRNA gene qPCR and amplicon sequencing indicated a negative effect of N on bacterial and archaeal abundance, species diversity, and richness. Our analyses revealed the predominance and cosmopolitan distribution of Proteobacteria, Bacilli, Anaerolineae, and Bacteroidota, their extensive adaptability and metabolic versatility under different N-amended conditions. Soil microbial community exhibited distinct responses to added N-substrates, with some bacterial taxa (Sphingomonadales) showing enhanced abundances while others were inhibited (Thermodesulfovibrionia and Clostridiales). The core-community across N-amended sets exhibited 68% taxonomic variation from its native counterpart, with two most abundant members of native soil, Nitrosomonadaceae and Beijerinckiaceae, diminishing, while Bacillaceae, Pseudomonadaceae, and Sphingomonadaceae became more prominent and resilient under N-amended conditions. The co-occurrence network highlighted that though total and core soil communities responded to N-amendments with an increased abundance of copiotroph bacteria, oligotrophs thrived under nutrient-limited conditions, serving as keystone taxa during prolonged fertilisation. These findings elucidate how synthetic N inputs restructure paddy soil microbiomes and alter their ecological functioning.

## Introduction

The community composition and function of the agricultural soil microbiome are critical components of the catabolic landscape of this ecosystem. These provide a repertoire of necessary biotic and abiotic resources to plants and other macroorganisms and maintain soil fertility and quality ^[1]^. The structure and dynamics of soil microbiome (bacteria and archaea) control the major biogeochemical cycle of soil and provide nutrients to promote plant growth ^[2, 3]^. With respect to understanding the soil microbiome for maintaining vital soil properties and fertility, paddy soil represents one of the most essential systems ^[4]^. Agricultural land used for the cultivation of rice (*Oryza*) covers 155 million hectares of the world’s surface and constitutes the largest anthropogenic wetland and agroecosystem, meeting the increasing demand of more than half of the global population’s caloric and essential micronutrient intake ^[5]^. The microbiome of the paddy soil has tremendous economic, social, and health significance and is under threat due to poorly managed and inefficient agriculture practices ^[6, 7]^. Inefficiency in soil management and various anthropogenic practices like application of excessive fertilizers, chemical pesticides and herbicides, chemical plant growth enhancers, and agrochemicals used to promote crop yield and injudicious exploitation of heavy metal contaminated groundwater for irrigation have disrupted soil microbial diversity and overall soil health, imposing nutritional imbalance and soil acidification, among other concerns ^[8–10]^. Other dangers include nutrient overflow, contamination, soil compaction, water pollution, and persistent toxicity ^[11–13]^. In light of the increased threat perceived, it is imperative to gain a better insight into the characteristics of soil microbiome, especially in response to its exposure to elevated fertilizer or other additives commonly practiced.

Among the various strategies adopted towards improving crop productivity, the addition of nitrogen (N) is noteworthy as enhanced N-supply allows meeting the crop and livestock production requirement for providing broader ecosystem services for local and global societies ^[14, 15]^. The global consumption of N has increased steadily to approximately 110 million metric tons in 2020 (from 105 million metric tons in 2010) ^[16, 17]^. Driven by the increased use of synthetic N, Southeast Asian countries, especially India and China, have gained self-sufficiency in crop production ^[18, 19]^However, imprudent water and N consumption for improved crop yield has inherited an unsustainable agricultural system ^[20, 21]^. Although various researchers have reported the ecological and environmental problems as footprints of embedded N overuse ^[4, 22]^, a comprehensive assessment of the impact (of N-amendments) on the tropical paddy soil microbial community in West Bengal is minimal, making it gain considerable scientific interest in due course of time. Various detrimental effects of excess and repeated N-amendment, deteriorating soil health and crop quality are reported in recent studies ^[23]^. It has been reported that global N addition reduces the overall soil microbial diversity by increasing competition for limited resources ^[24–26]^. For example, some specific soil microorganisms, such as those that prefer acid and ammonium (NH_4_^+^), benefit from N enrichment in N-limited conditions ^[27]^. Under N-enriched conditions, however, the intense competition causes these microbial species to essentially die out, reducing the overall diversity of the soil microbial community ^[28]^.

Over the past few years, there has been rigorous research on agricultural practices, particularly those involving extensive use of synthetic fertilizers, pesticides and herbicides, intensive tillage, chemical plant growth enhancers, and heavy metal contamination due to anthropogenic activities that have been shown to impact soil microbiomes negatively. In India, Kumar et al. ^[29, 30]^, observed that N-application on farm soil (developed from deltaic sediments and classified as Aeric Endoaquept, with a sandy clay loam texture) alone suppressed several beneficial bacterial phyla (Proteobacteria, Acidobacteria, Cyanobacteria, Fibrobacteres, Spirochaetes, TM7, and GNO4) and a significant reduction in diazotrophic taxa, resulting in the alteration of soil biodiversity and rice productivity. But despite these findings, there is a lack of studies focusing on Indian paddy soils using microcosm-based experiments and 16S rRNA amplicon sequencing to understand the immediate effects of N addition on soil microbial communities and characterize the changes in microbial communities from N-amendment. Addressing this gap is necessary to obtain valuable insights into sustainable agricultural practices and soil health management in Indian paddy ecosystems. Ecological research in India has focused on measuring microbial communities and some key soil functions under long-term N application, making the initial response of the microbial communities remain unclear ^[29–31]^. Thus, the impact of excess N application on soil microbial communities remains a significant knowledge gap in agricultural soil microbiome research.

The present study aimed to gain deeper insights into the impact of various N-amendments on paddy soil microbiome by assessing the microbial cell abundance and response of the microbiome, allowing enrichment of specific populations under different N-amendments. The primary questions addressed through this study are: (i) How does the soil bacterial community respond to the amendment of different N-compounds, and (ii) What are the unique bacterial populations enriched in response to different N-compounds. This understanding of the structure and function of soil microbial communities involved with N mineralization change in response to N-amendment practices provides critical information on the effect of enhanced N-compound addition to paddy soil microbial ecology, which would be essential for developing sustainable agricultural practices by reducing the negative agroecology consequences.

## Materials and methods

### Soil sampling and characterization

Soil samples used in this study were collected from agricultural fields of Bethuadahari, Nadia, one of the arsenic (As) hotspots in West Bengal ^[32]^. Samples were collected 10–20 cm below the soil surface from ten agricultural sites. Three to five subsamples from each site were collected, mixed thoroughly, and stored in sterile containers. The samples were immediately stored at 4°C until shipped to the laboratory. Elemental composition and various other physicochemical properties of the soil samples were determined and reported in a previous study ^[32]^.

### Microcosm experiments

Based on the geographic locations, soil samples were mixed into two soil mixtures (S1 and S2) for further microcosm-based experimental study. To set up the microcosms, 10 g (wet-weight) soil was added to a 120 mL serum vial that contained 60 mL of Tris-buffered minimal salt medium ^[33]^. A cocktail of carbon substrates (5 mM each of sodium carbonate, glucose, sodium gluconate, sodium lactate, and sodium acetate) was added along with specific N substrates. The initial pH of the microcosm was set at 7.2 ± 0.2. Four N-amendments were made to study the effect of different N-substrates, viz, nitrite (NO_2_^−^), nitrate (NO_3_^−^), ammonium (NH_4_^+^), and urea (CO(NH_2_)_2_) (each 5mM). A no N-amended set was maintained as control (No N). All the microcosms were prepared in duplicates and incubated at 25 °C for three hundred days. Details of the microcosms and their physicochemical details are presented in Table 1.

**Table 1:** Details of the microcosm setups and their physicochemical properties.

| Sl. no. | Microcosm ID | Treatment | Brief Description | pH | EC (mS/cm) | DO (mg/L) | ORP (mV) |
| --- | --- | --- | --- | --- | --- | --- | --- |
| 1 | S1_1 | No N | No N added | 7.5 ± 0.08 | 7.75 ± 0.09 | 1.88 ± 0.15 | -32.7 ± 0.06 |
| 2 | S1_2 | NO<sub>2</sub><sup>-</sup> | Soil amended with NO<sub>2</sub><sup>-</sup> nitrogen (5mM) | 6.6 ± 0.10 | 8.36 ± 0.20 | 0.90 ± 0.11 | -133.4 ± 0.97 |
| 3 | S1_3 | NO<sub>3</sub><sup>-</sup> | Soil amended with NO<sub>3</sub><sup>-</sup> nitrogen (5mM) | 6.5 ± 0.08 | 8.18 ± 0.08 | 0.88 ± 0.09 | -142.7 ± 0.84 |
| 4 | S1_4 | NH<sub>4</sub><sup>+</sup> | Soil amended with NH<sub>4</sub><sup>+</sup> nitrogen (5mM) | 6.5 ± 0.09 | 7.77 ± 0.24 | 0.62 ± 0.09 | -160.2 ± 0.52 |
| 5 | S1_5 | Urea | Soil amended with Urea nitrogen (5mM) | 6.8 ± 0.12 | 7.61 ± 0.12 | 0.69 ± 0.06 | -168.8 ± 0.62 |
| 6 | S2_1 | No N | No N added | 6.2 ± 0.09 | 7.87 ± 0.09 | 1.25 ± 0.21 | -57.8 ± 0.38 |
| 7 | S2_2 | NO<sub>2</sub><sup>-</sup> | Soil amended with NO<sub>2</sub><sup>-</sup> nitrogen (5mM) | 6.4 ± 0.19 | 7.65 ± 0.15 | 1.18 ± 0.15 | -50.6 ± 0.46 |
| 8 | S2_3 | NO<sub>3</sub><sup>-</sup> | Soil amended with NO<sub>3</sub><sup>-</sup> nitrogen (5mM) | 6.4 ± 0.09 | 7.95 ± 0.34 | 0.31 ± 0.13 | -134.3 ± 0.94 |
| 9 | S2_4 | NH<sub>4</sub><sup>+</sup> | Soil amended with NH<sub>4</sub><sup>+</sup> nitrogen (5mM) | 6.4 ± 0.09 | 8.13 ± 0.16 | 0.76 ± 0.09 | -123.2 ± 0.41 |
| 10 | S2_5 | Urea | Soil amended with Urea nitrogen (5mM) | 6.6 ± 0.08 | 8.13 ± 0.24 | 0.51 ± 0.08 | -131.8 ± 0.85 |

### Physicochemical analyses

Dissolved oxygen, oxidation-reduction potential, pH, and conductivity of the microcosm setups were measured using highly sensitive probes fitted with an Orion Multiparameter (Thermo Fisher Scientific). The soluble fraction was extracted from the microcosm through centrifugation (10000g, 15min), and ions such as SO_4_^2−^, PO_4_^3−^, NO_2_^−^, NO_3_^−^, NH_4_^+^, As^5+^, Na^+^, Mg^2+^, Ca^2+^, and K^+^ were estimated from this through Ion Chromatography (Eco IC, Metrohm) ^[34]^. The micro-Kjeldahl method estimated the total N (of the soil) and aqueous N as per the procedure suggested by AOAC ^[35]^. Elemental quantitation (density scanning) of the sediments from the microcosm was performed using a field emission-scanning electron microscope (FE-SEM, ZEISS EVO) coupled with an X-ray spectrometer (EDS, ZEISS). The aqueous phases Fe and As were measured using US EPA 6010D methods through Inductively Coupled Plasma-Optical Emission Spectrometry (ICP-OES). All analyses were performed in triplicate.

### DNA extraction and 16S rRNA gene amplification and quantification through qPCR

The total community DNA from each microcosm sample was extracted in duplicate (using 5 mL slurry) using E.Z.N.A.® Soil DNA Kit (Omega Bio-tek) following the manufacturer’s instructions and pooled. For DNA quantification, 1 μl of each sample was used for determining concentration using Qubit® 3.0 Fluorometer (Thermo Fisher). The gene copy numbers of the 16S rRNA bacterial gene were determined by quantitative PCR (QuantStudio 5, Thermo Fisher) by using SYBR green as the detection system (Power Syber Green PCR Master Mix) ^[36]^. Thermal cycling, fluorescent data collection, and data analysis were carried out with the QuantStudio 5 sequence detection system according to the manufacturer’s instructions.

### 16S rRNA gene library preparation and high-throughput sequencing

Post quantification, total DNA samples were subjected to Amplicon PCR, which was set up using V3V4 primers (IDT) and Agilent Paq5000 2X Master Mix. The DNA was diluted to 1ng/μL for PCR setup, while the primer concentration was 10μM. The desired PCR product size was ~450bp. 1X bead-based clean-up was performed post-Amplicon PCR using Kapa beads. The thermal cycler used for library preparation was Agilent-Sure Cycler. “Kapa Hyper Prep Kit” from Roche was used for library preparation, followed by further processing, and subjected to sequencing on Illumina Miseq.

### Bioinformatic analysis of 16S rRNA gene sequence data

Demultiplexed paired-end reads were assembled according to the expected amplicon length (430 - 470 bp for 16S rRNA gene) and filtered by maximum error rates of 0.5% using USEARCH v9.1.13 (32-bit) ^[37]^. Potential chimeric sequences were detected and removed in QIIME 2 ^[38]^ using VSEARCH. Then, the resulting gene sequences were taxonomically classified into operational taxonomic units (OTUs) with a 0.03 dissimilarity cut-off using QIIME. Yet, singletons were excluded since they likely originated from sequencing errors. Sequences of all samples were normalized to the same number (i.e., the lowest number of sequences of all samples, 11403 for the 16S rRNA gene) to make better and more reliable comparisons of microbial diversity among different samples. For 16S rRNA gene amplicons, OTUs were picked up with open reference of the 16S SILVA 138 reference database (https://www.arb-silva.de/documentation/release-138) using the method of USEARCH v6.1.544 (Beta). OTUs were analyzed for alpha diversity, including richness (observed OTUs, Chao1), evenness, and Shannon diversity index.

### Statistical analyses

Principal Coordinates Analysis (PCA) was done with PAST v4.12b, and significant changes in the microbial assemblages and other metrics generated from various N-amendments were examined (https://palaeo-electronica.org). The Bray–Curtis similarity index was used to assess the statistical significance of the similarity of various N-amendments across samples. Similarity Percentage (SIMPER) analysis was done with PAST v4.12b to identify the taxa contributing to the average dissimilarity between control and N-treated conditions. The core community was represented in a Venn diagram using EVenn ^[39]^. Co-occurrence networks were constructed using the lowest taxonomy level of the core community. The Spearman correlations (pairwise) among the taxa were calculated for co-occurrence networks, and only the strong correlations (r ≥ ± 0.7; p < 0.01) were considered for visualization of interaction network using Cytoscape 3.10.2 ^[40]^. All plots were made using OriginPro 2023 and R Studio 4.3.2 (http://www.r-project.org). For 16S rRNA gene sequences, OTUs abundant in the taxa that were most affected by N-amendments and that remained unclassified at various taxonomic levels were searched for similarity using the curated species database of EzTaxon Biocloud server (https://www.ezbiocloud.net/) and NCBI database (https://blast.ncbi.nlm.nih.gov/) followed by molecular phylogenetic analysis by Neighbour Joining method based on Jukes-Cantor model in MEGA 11 ^[41]^ and visualization through iTOL (https://itol.embl.de/) ^[42]^. Native soil microbial community was reanalyzed for the purpose of studying the effect of the N-amendment on core communities (before and after the N-amendment). 16S rRNA gene amplicon sequence data previously obtained from the native soils, available in the NCBI (BioProject Number: PRJNA686650) database ^[32]^, was reanalyzed for core-community of native soil as a part of this study. 16S rRNA gene amplicon sequence data retrieved from the microcosms and used in this study are deposited in the NCBI Database under the BioProject Number: PRJNA1000929, accession numbers: SRR25479801-SRR25479810.

## Results

### Soil and microcosm characteristics

The physicochemical characteristics of the N-amended microcosms are presented in Table 1. All the treatments exhibited reducing conditions (ORP: −168.8mV to −32.7mV), increased dissolved ions (indicated by conductivity), and low dissolved oxygen (0.31-1.88 mgL^−1^). Analysis of the aqueous phase showed that N-amendment (maximum in NH_4_^+^ and urea amended sets) increased the concentrations of soluble Ca^2+^, Mg^2+^, SO_4_^2−^, and total (soluble) Fe across the microcosms (Fig. 1a). Principal component analysis confirmed NH_4_^+^ and urea amendment facilitated release of major soil ions (Na^+^, Ca^2+^, Mg^2+^, total soluble Fe and soluble SO_4_^2−^) (Fig. 1b). Elemental mapping, done through the energy-dispersive X-ray (EDX) analysis of the solid phase, revealed the presence of N, Al, Si, and Fe (Fig. 1c). A notable alteration in the N concentrations was observed after incubation (Fig. S1). Microscopic investigation revealed the presence of mixed culture in the N-amended sets (Fig. S2).

**Figure 1.**
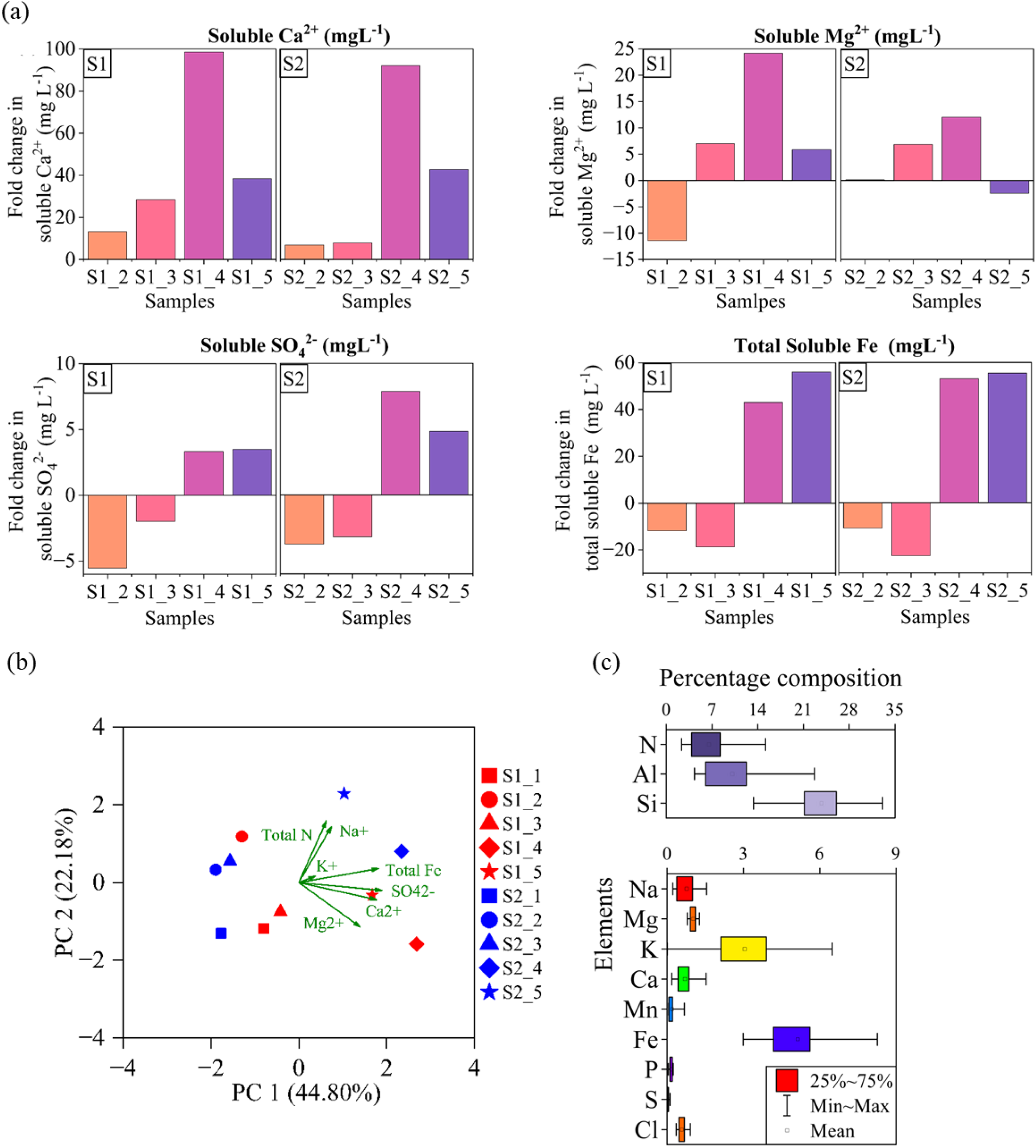
(a) Fold change in concentrations of Ca^2+^, Mg^2+^, SO_4_^2−^ and total Fe following incubation, (b) Principal Component Analysis (PCA) based on major chemical parameters. The variables were scaled prior to the analysis, and (c) Major elemental composition of the solid phases as measured through EDX spectral scanning

### Quantitative estimation of bacterial and archaeal abundance and alpha diversity analysis from 16S rRNA gene amplicon sequencing data

Quantitative assessment of bacterial abundance in the soil microcosms portrayed distinct effects of the amendment of various N sources (Table 2). Compared to the native soil ^[32]^, bacterial and archaeal 16S rRNA gene copies were increased within the no-N-amended sets (S1_1 and S2_1). Noticeably, among all the microcosm sets, the maximum number of 16S rRNA gene copies were observed in these two sets (S1_1 and S2_1; 16.6 × 10^8^ and 15.6 × 10^8^ for bacteria, and 38.3 × 10^5^ and 26.1 × 10^5^ for archaea, respectively), which did not receive any additional N supplementation. The extent of enhancement in cell number varied among the two soil mixtures tested (S1 and S2). In contrast, all the N-amended microcosms showed relatively lower 16S rRNA gene copy numbers (Table 2).

**Table 2:**
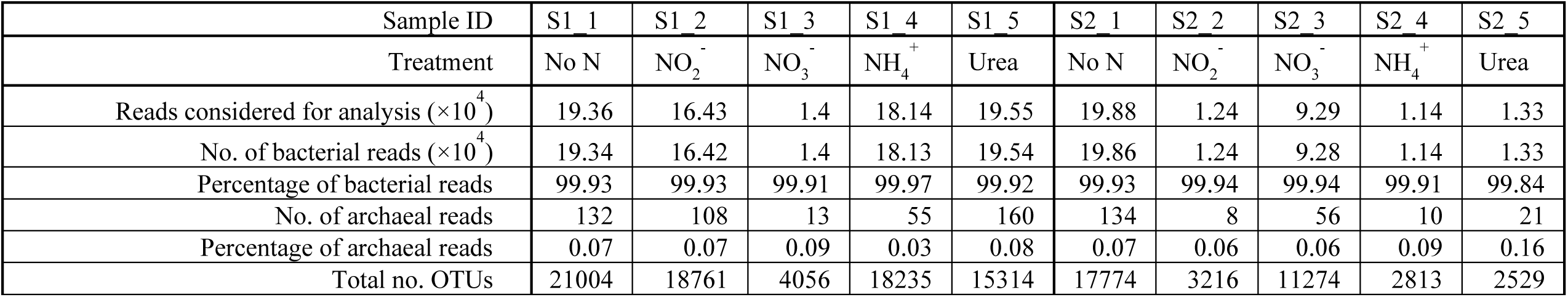
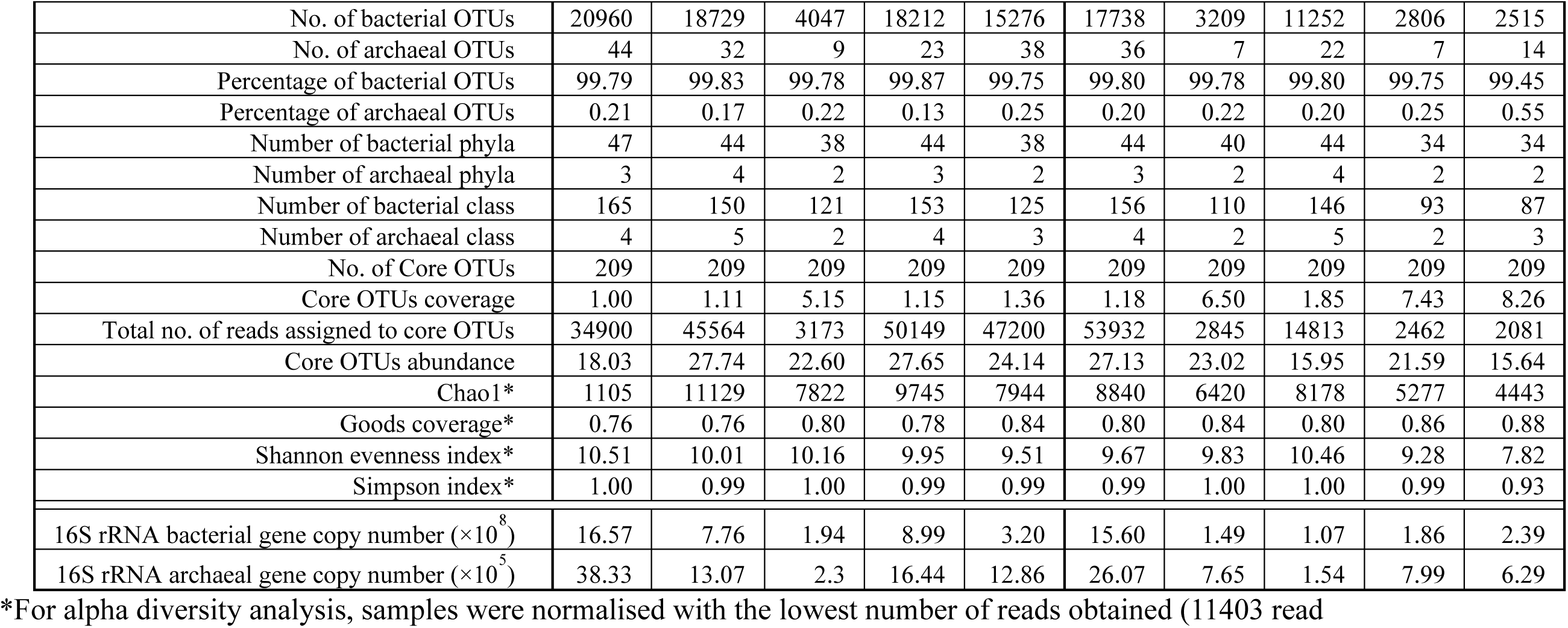
Sequencing read details, alpha diversity parameters and qPCR-based abundances of 16S rRNA genes.

| Sample ID | S1_1 | S1_2 | S1_3 | S1_4 | S1_5 | S2_1 | S2_2 | S2_3 | S2_4 | S2_5 |
| --- | --- | --- | --- | --- | --- | --- | --- | --- | --- | --- |
| Treatment | No N | NO<sub>2</sub><sup>-</sup> | NO<sub>3</sub><sup>-</sup> | NH<sub>4</sub><sup>+</sup> | Urea | No N | NO<sub>2</sub><sup>-</sup> | NO<sub>3</sub><sup>-</sup> | NH<sub>4</sub><sup>+</sup> | Urea |
| Reads considered for analysis (×10<sup>4</sup>) | 19.36 | 16.43 | 1.4 | 18.14 | 19.55 | 19.88 | 1.24 | 9.29 | 1.14 | 1.33 |
| No. of bacterial reads (×10<sup>4</sup>) | 19.34 | 16.42 | 1.4 | 18.13 | 19.54 | 19.86 | 1.24 | 9.28 | 1.14 | 1.33 |
| Percentage of bacterial reads | 99.93 | 99.93 | 99.91 | 99.97 | 99.92 | 99.93 | 99.94 | 99.94 | 99.91 | 99.84 |
| No. of archaeal reads | 132 | 108 | 13 | 55 | 160 | 134 | 8 | 56 | 10 | 21 |
| Percentage of archaeal reads | 0.07 | 0.07 | 0.09 | 0.03 | 0.08 | 0.07 | 0.06 | 0.06 | 0.09 | 0.16 |
| Total no. OTUs | 21004 | 18761 | 4056 | 18235 | 15314 | 17774 | 3216 | 11274 | 2813 | 2529 |
| No. of bacterial OTUs | 20960 | 18729 | 4047 | 18212 | 15276 | 17738 | 3209 | 11252 | 2806 | 2515 |
| No. of archaeal OTUs | 44 | 32 | 9 | 23 | 38 | 36 | 7 | 22 | 7 | 14 |
| Percentage of bacterial OTUs | 99.79 | 99.83 | 99.78 | 99.87 | 99.75 | 99.80 | 99.78 | 99.80 | 99.75 | 99.45 |
| Percentage of archaeal OTUs | 0.21 | 0.17 | 0.22 | 0.13 | 0.25 | 0.20 | 0.22 | 0.20 | 0.25 | 0.55 |
| Number of bacterial phyla | 47 | 44 | 38 | 44 | 38 | 44 | 40 | 44 | 34 | 34 |
| Number of archaeal phyla | 3 | 4 | 2 | 3 | 2 | 3 | 2 | 4 | 2 | 2 |
| Number of bacterial class | 165 | 150 | 121 | 153 | 125 | 156 | 110 | 146 | 93 | 87 |
| Number of archaeal class | 4 | 5 | 2 | 4 | 3 | 4 | 2 | 5 | 2 | 3 |
| No. of Core OTUs | 209 | 209 | 209 | 209 | 209 | 209 | 209 | 209 | 209 | 209 |
| Core OTUs coverage | 1.00 | 1.11 | 5.15 | 1.15 | 1.36 | 1.18 | 6.50 | 1.85 | 7.43 | 8.26 |
| Total no. of reads assigned to core OTUs | 34900 | 45564 | 3173 | 50149 | 47200 | 53932 | 2845 | 14813 | 2462 | 2081 |
| Core OTUs abundance | 18.03 | 27.74 | 22.60 | 27.65 | 24.14 | 27.13 | 23.02 | 15.95 | 21.59 | 15.64 |
| Chao1* | 1105 | 11129 | 7822 | 9745 | 7944 | 8840 | 6420 | 8178 | 5277 | 4443 |
| Goods coverage* | 0.76 | 0.76 | 0.80 | 0.78 | 0.84 | 0.80 | 0.84 | 0.80 | 0.86 | 0.88 |
| Shannon evenness index* | 10.51 | 10.01 | 10.16 | 9.95 | 9.51 | 9.67 | 9.83 | 10.46 | 9.28 | 7.82 |
| Simpson index* | 1.00 | 0.99 | 1.00 | 0.99 | 0.99 | 0.99 | 1.00 | 1.00 | 0.99 | 0.93 |
| 16S rRNA bacterial gene copy number ($\times 10^8$) | 16.57 | 7.76 | 1.94 | 8.99 | 3.20 | 15.60 | 1.49 | 1.07 | 1.86 | 2.39 |
| 16S rRNA archaeal gene copy number ($\times 10^5$) | 38.33 | 13.07 | 2.3 | 16.44 | 12.86 | 26.07 | 7.65 | 1.54 | 7.99 | 6.29 |

\*For alpha diversity analysis, samples were normalised with the lowest number of reads obtained (11403 read)

The 16S rRNA gene-based amplicon sequencing yielded a considerable number of high-quality reads (4.9 × 10^5^ reads, on average per sample) from each sample. Following quality check, including the removal of singletons, a total of 10.7 × 10^5^ reads in total were obtained and were clustered (99% identity threshold) into 114976 Operational Taxonomic Units (OTUs) (Table 2). On average, 99.7% of these OTUs were associated with bacteria, while archaea constituted a minute fraction (average 0.2 %). The OTUs’ taxonomic association revealed 2-4 archaeal and up to 47 bacterial phyla (Table 2).

Alpha diversity indices (Chao1, Simpson, and Shannon) were calculated to assess microbial community abundance and consistency (Table 2). The no N-amendment set exhibited the highest species diversity and abundance, while the N-amendment decreased diversity and abundance in all microcosms. This decline and reduced 16S rRNA gene copy numbers from qPCR data indicated a gross negative impact of N-amendment on soil microbial communities (Table 2). Notably, 209 OTUs (1.0 - 8.3% of the total 114976 OTUs) were found to be shared (100% prevalence) across all the ten samples, representing the core community resilient to varied N sources. These core OTUs comprised 15.6 - 27.7% of each community.

### Microbial community structure and community response towards N-amendments

Taxonomic analysis of the communities showed an overall predominance of Gamma- and Alphaproteobacteria, Bacilli, Bacteroidota, Anaerolineae, and Acidobacteriota (Fig. 2a). These dominant taxa demonstrated their ubiquitous presence (with varied abundances) across all the sets. Gemmatimonadota, Myxococcota, Actinobacteria, Thermoleophilia, Verrucomicrobiota, Dehalococcoidia, and Desulfobacterota were detected as other abundant taxa. These were followed by some other members, namely, Planctomycetota, Patescibacteria, Armatimonadota, Clostridia, Chloroflexi_KD4-96, Nitrospirota, and Cyanobacteria with considerably lower abundances (Table S3).

**Figure 2.**
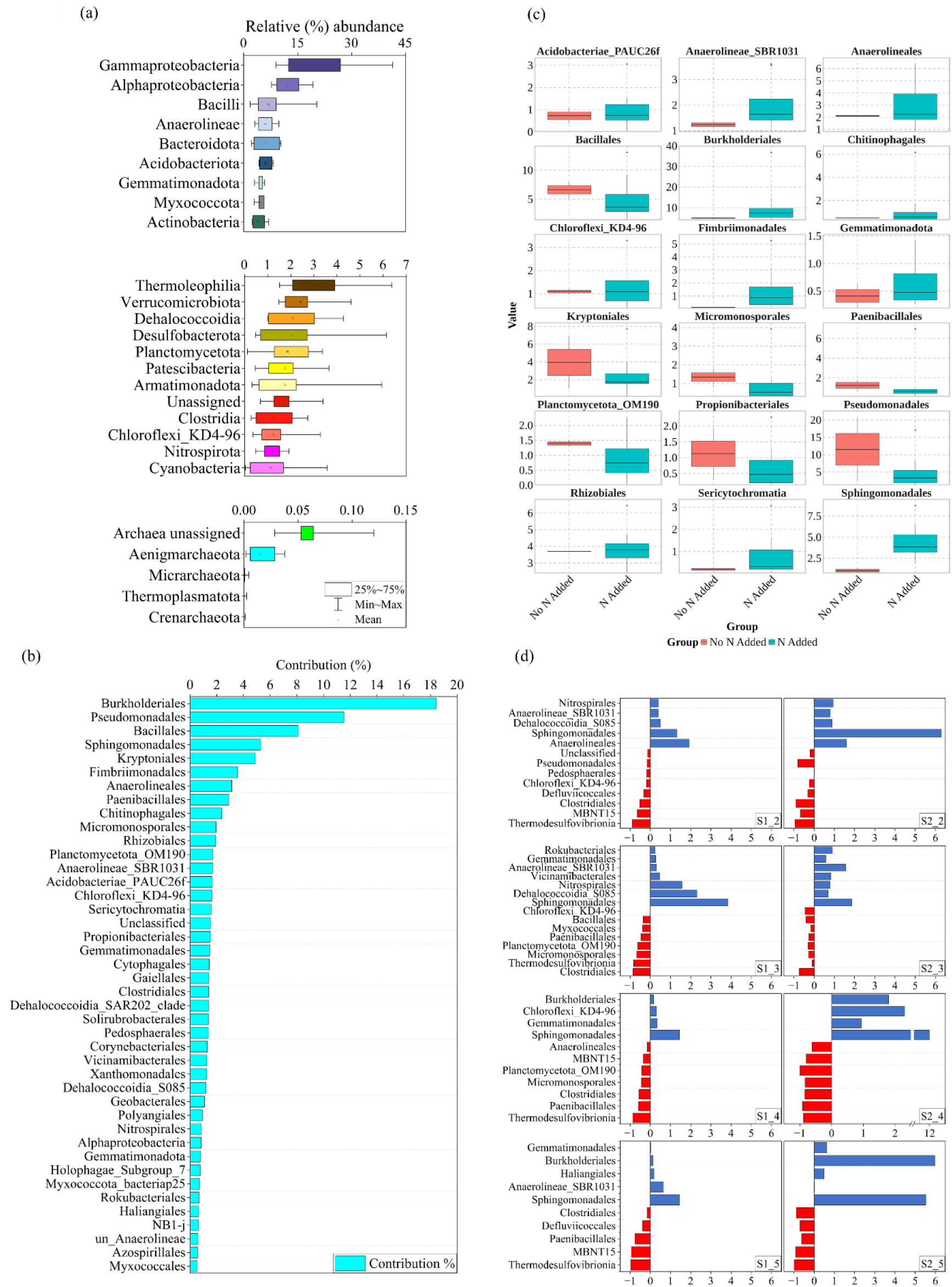
(a) Overall microbial community composition (Phylum-Class level) across the samples. Taxa having average abundance > 1% in the total community were only considered, (b) Similarity percentage (SIMPER) analysis displaying the contribution of individual taxa (order level) in explaining the difference between no N added and N added communities, (c) Distribution of the bacterial taxa contributing to the major differences (contribution >1.5%) between no N added and N added conditions (taxa were selected based on the SIMPER analysis), and (d) Fold change (with respect to control, No N added) of the shared taxa between S1 and S2-communities affected negatively and positively due to different N amendment

Similarity percentage (SIMPER) analysis was performed between No N and N-amended sets to identify the bacterial taxa that were most affected (either positively or negatively) during the N-amendment (Fig. 2b). As evidenced, members of Burkholderia, Pseudomonadales, Bacillales, Sphingomonadales, Kryptoniales, Fimbriimonadales, Anaerolineales and a few more taxa were found to be most affected. Considering all the N-amended sets in one group and no N-amended sets in another, an overall analysis revealed that Burkholderia, Sphingomonadales, Fimbriimonadales, Anaerolineae_SBR1031, Anaerolineales, Gemmatimonadota and Chitinophagales were more positively impacted due to the amendment of N-amendments, while Kryptoniales, Bacillales, Acidobacteria_PAUC26f, Micromonosporales, Paenibacillales, Pseudomonadales, and Propionobacteriales were negatively impacted (Fig. 2c).

The individual effect of different N-amendments on the microbial community was compared in both soils (Fig. 2d). The addition of NO_2_^−^ had promoted the growth of Sphingomonadales, Anaerolineales, Dehalococcoidia_S085, Anaerolineae_SBR1031, and Nitrospirales. Among the taxa that positively responded to NO_2_^−^ many also responded favorably in the presence of NO_3_^−^. Additionally, Rokubacteriales, Vicinamibacterales, and Gemmatimonadales were also preferred by NO_3_^−^. Sphingomonadales, Burkholderiales, Chloroflexi_KD4-96 and Gemmatimonadales; and Anaerolineae_SBR1031, and Haliangiales showed a favorable response towards NH_4_^+^ and urea amendments. Noticeably, members of obligate anaerobic taxa like Thermodesulfovibrionia, Clostridiales, and Chloroflexi_KD4-96 were negatively impacted by both NO_2_^−^ and/or NO_3_^−^amendment. A few more taxa were discretely inhibited as well, *e.g.*, MBNT15, Defluviicoccales, Pedosphaerales, and Pseudomonadales in NO_2_^−^; and Planctomycetota_OM190, Paenibacillales, Myxococcales, Bacillales and Micromonosporales in NO_3_^−^. MBNT15, Paenibacillales, Anaerolineales Micromonosporales, Planctomycetota_OM190, Defluviicoccales, Thermodesulfovibrionia and Clostridiales taxa were negatively impacted by either NH_4_^+^ or Urea or both. Thus, microbial response to various N substrates depicted that Sphingomonadales increased in both soils regardless of the N substrate, followed by Nitrospirales, Dehalococcoidia_S085, Anaerolineae_SBR1031, Gemmatimonadales, and Burkholderiales, which majorly showed an increased abundance. In contrast, taxa such as Kryptoniales, Gaiellales, and others varied in their response between the two soils. Notably, Thermodesulfovibrionia, Clostridiales, and a few others exhibited a significant decrease in abundance across both soils, regardless of the N-amendments.

### Core Community

A further analysis was performed to identify the distribution of core/endemic community, which focused on 209 OTUs shared across all 10 microcosm sets (including the No N amended set) (Table 2, Fig. 3a). These OTUs, representing 1-8.26% of total OTUs per sample, accounted for a significant total abundance of 15.64–27.74%. Taxonomic affiliation of these core OTUs indicated the dominance of significant phyla such as Gamma and Alpha-proteobacteria, Bacilli, Gemmatimonadota, Thermoleophilia, Dehalococcoidia and others (Fig. 3b). At the genus level, 93 taxa were identified, with Pseudomonas, Bacillus, Sphingomonas, Gemmatimonadaceae, and Ammoniphilus being the most prevalent (Fig. S3).

**Figure 3.**
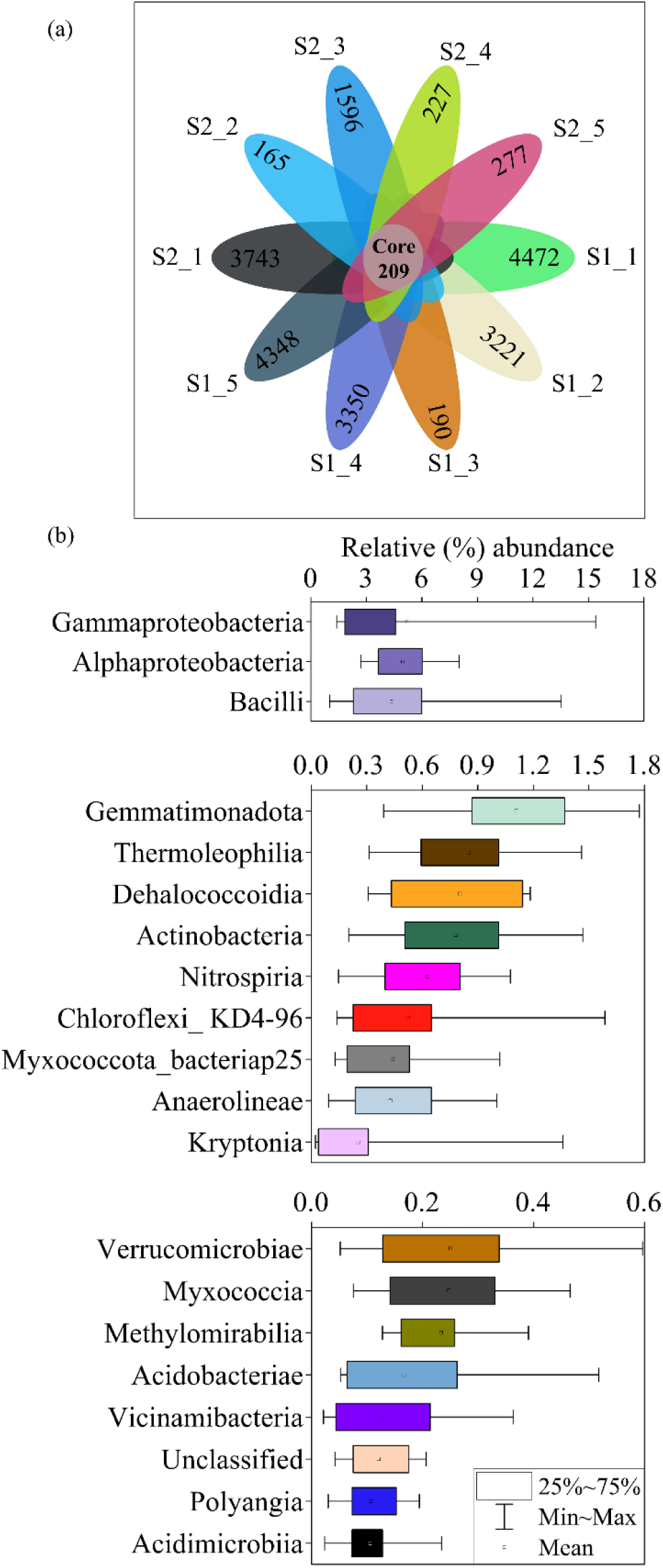
(a) Venn diagram displaying Operational Taxonomic Units (OTUs) detected in all the ten communities. The numbers displayed on the leaves of the Venn represented the number of unique OTUs detected in that sample. The total numbers of OTUs shared among all the samples are displayed in the centre and considered core OTUs. (b) There is a relative abundance of core community at the Phylum Class level. Taxa having average abundance > 0.1% in the core community were considered, and (c) Canonical Correspondence analysis (CCA) based on core community members and major physiochemical parameters of the N-microcosms

To understand the effect of the N-amendment on the core community of paddy soil, the native soil core community was reanalyzed together with N-amended setups. It was observed that 261 Core OTUs were present in the native soil. However, among the 10 microcosm setups, 8 were N-amended sets, which shared 227 endemic OTUs. In native soil, these OTUs belonged to 78 families, while N-treated conditions had 89 diverse families, revealing 68% taxonomic variation with the native soil. The top 20 families showed that Nitrosomonadaceae, Beijerinckiaceae, Sphingomonadaceae, Burkholderiaceae, and Nitrososphaeraceae dominated in native soil (Fig. 4a), which either diminished or perished under N-amended conditions, while Bacillaceae, Pseudomonadaceae, Sphingomonadaceae, Gemmatimonadaceae, and Paenibacillaceae were more prevalent in N-amendment (Fig. 4b).

**Figure 4.**
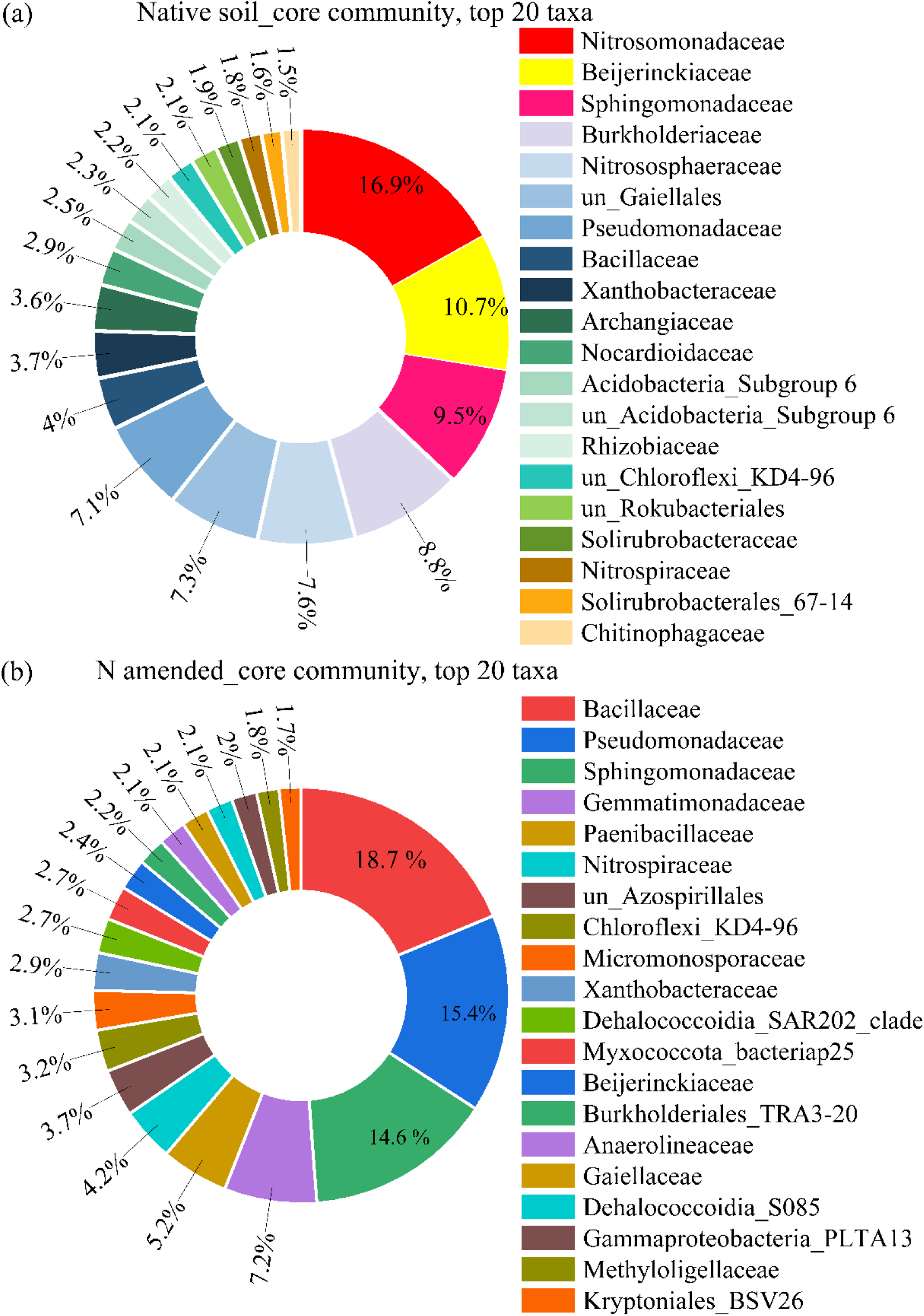
Comparison between the core communities (family level) between (a) native soil and (b) N-treated soil

### Co-occurrence network analysis

Co-occurrence patterns among the core community members (affiliated with 93 Core OTUs) were inferred based on strong and significant correlations (r > ± 0.7, p < 0.01) to understand the microbial community interactions (Fig. 5). The positive network (r > 0.7) comprised 71 nodes and 153 edges, with an average shortest path length of 4.319, network density of 0.064, and clustering coefficient of 0.361 (Table S4). Based on the degrees of connection, a number of keystone families were identified. These families were affiliated with unclassified Alphaproteobacteria, Candidatus_Udaeobacter, and Azospirillales, which had the highest degree values (Fig. 5a). The negative network (r < −0.7) had 17 nodes and 16 edges, with an average shortest path length of 2.167 and network density of 0.197 (Table S4). Dominant taxa included Dadabacteriales and Methyloligellaceae (Fig. 5b). The network was divided into 7 modules, indicating robust intra-module connectivity and cooperation among taxa (Fig. S3).

**Figure 5.**
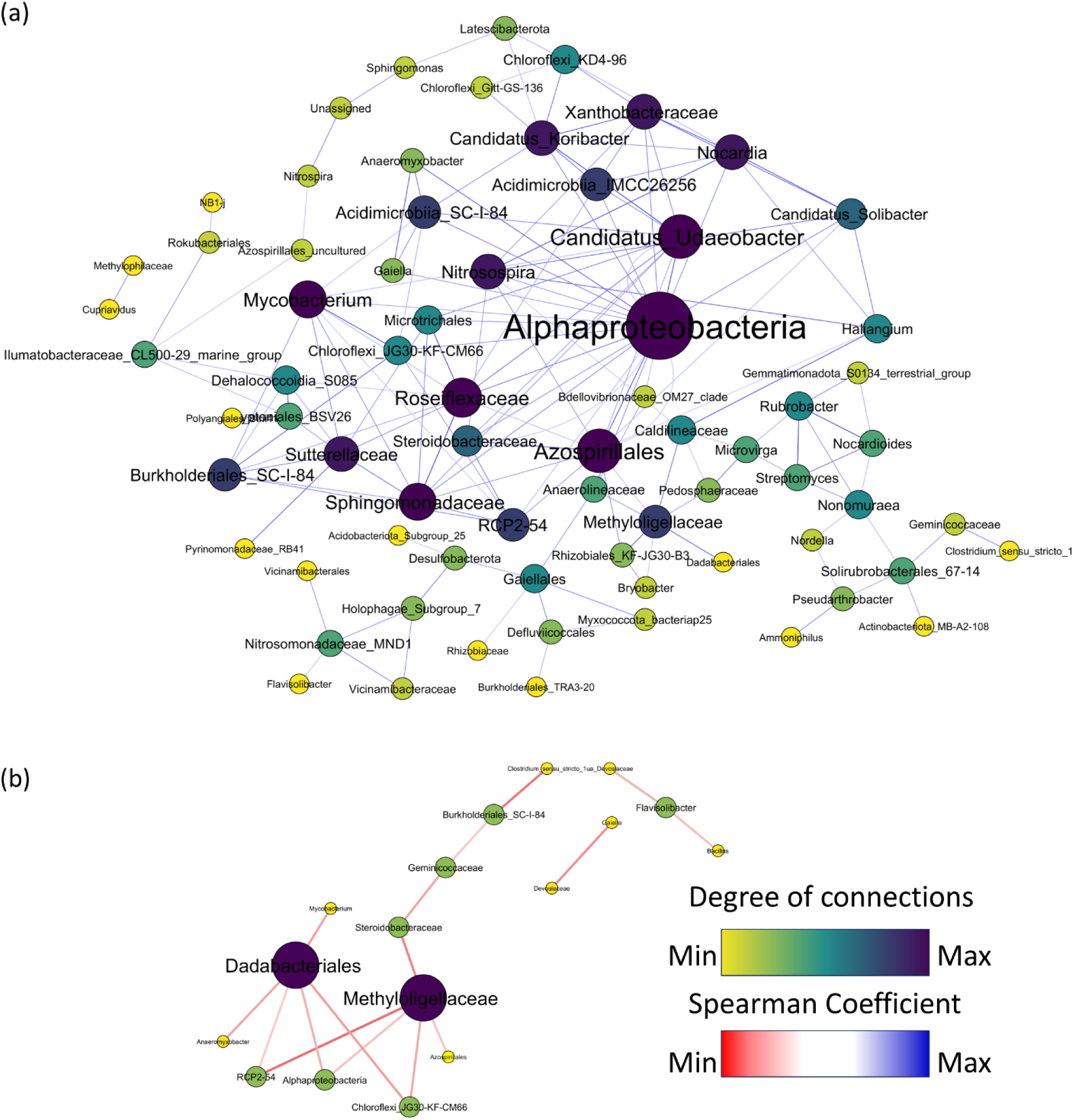
Co-occurrence network of the core community (lowest taxonomy level). Node size and color are proportional to the number of degrees. Nodes with higher degrees are bigger in size and purple in color than those with lower connections. The color of the edges represents the strength of Spearman’s correlations between the nodes. The strength of the connections is indicated in blue as being stronger and red as being weaker. For the Spearman correlation values, |r| > 0.7 are only considered, (a) indicating positively correlated taxa, and (b) representing negatively correlated taxa

### Phylogenetic analysis

Major OTUs, which are unclassified at the lowest taxonomy levels and were broadly affiliated to the taxa that were most affected by the N-amendment, regardless of the N substrate, were considered for analyzing the phylogeny. This analysis was performed using the most similar sequences available in the public database. These OTUs from different N-amended microcosms (members of the top 5 OTUs of each microcosm) represented the communities phylogenetic analysis suggested taxonomic affiliation of these OTUs to diverse orders of Sphingomonadales, Dehalococcoidia, Burkholderiales, Thermodesulfovibrionia and Clostridiales (Fig. 6), which belonged to the diverse sites well studied for N metabolism.

**Figure 6.**
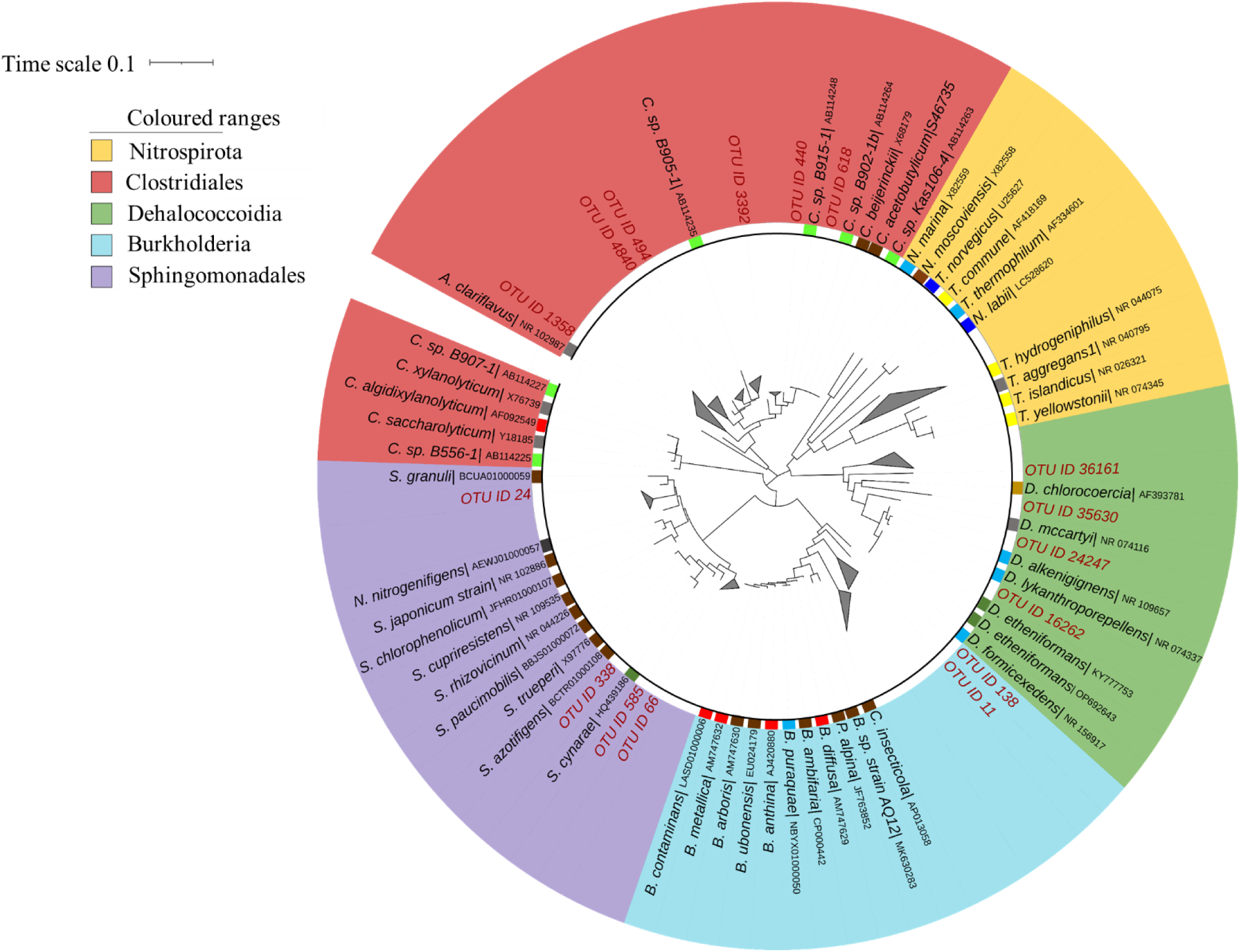
Phylogenetic analysis of the major OTUs affected (positively and negatively following N-amendments) due to N-amendment. The relative abundance of these OTUs across the samples is provided in Supplementary File 1

## Discussion

Microorganisms in paddy soil have been reported to be vital for material circulation, energy flow, nutrient transformation, and organic matter decomposition, serving as soil health indicators ^[1, 43]^. However, synthetic nitrogen was previously believed to impact these microbial communities in paddy soil ^[23, 44]^. This study employed a culture-dependent microcosm approach to evaluate the effects of different nitrogen amendments on paddy soil microbial communities.

### Influence of N on soil physicochemical factors

The application of N-amendments was previously reported to affect the mobilization of soil ions ^[45–47]^. A similar observation was noted in this study, where N-amendments, particularly NH_4_^+^ and urea, significantly increased the concentrations of various ions, including Na^+^, Ca^2+^, Mg^2+^, SO_4_^2−^, and total Fe in the aqueous phase, altering the gross soil physicochemical properties. The principal component analysis corroborated that majorly NH_4_^+^ and urea amendment aided the release of these soil ions. Elevated N levels have been reported to influence soil chemistry and mineral stability through processes like nitrification and ferrolysis, which involve the reduction of iron (III) oxides to Fe^2+^ under wet conditions, leading to the release of cations and the destruction of clay minerals ^[48, 49]^.

Thus, enhanced N availability might have affected the redox conditions and pH, facilitating the mobilization of elements and ions. The shifts in these physicochemical parameters have been reported to facilitate mineral-microbe interactions and nutrient solubilization ^[50–52]^. Hao et al., 2020 ^[53]^ observed that increased soil acidification from N application decreased pH and exchangeable base cations, primarily due to higher nitrification rates. Urea hydrolysis might have initially raised soil pH through NH_4_ ^+^ production, but subsequent nitrification led to acidification. Thus, these analyses supported previous findings on N’s role in modifying soil chemical properties and enhancing nutrient availability.

### Effect of N-amendment on Bacterial 16S rRNA Gene abundance and α-diversity

This study observed that N-amendment decreased the 16S rRNA copy number in soil, indicating reduced bacterial and archaeal abundance due to lower growth rate in higher N concentrations. Previously, it was reported that interactions between various N-amendments in the soil ecosystem affected soil microbiome, microbial processes, and their abundance ^[54–56]^. The observed reduction in bacterial and archaeal 16S rRNA gene copy number might be attributed to the alterations in overall nutrient availability and altered patterns of interactions among the soil microbial populations in response to N-amendments, corroborated by the change in the physicochemical parameters. Together with the qPCR data (showing a reduction in 16S rRNA gene copy number), the decline in species diversity as observed in alpha diversity analysis (of 16S rRNA gene amplicons) represented a gross negative impact of N-amendment on the soil microbial community (with the highest bacterial diversity index observed with no N-amended control). Low to moderate levels (< 150 N kg ha^−1^ y^−1^) of available N have been reported to enhance diversity, while high levels suppress the same (> 150-300 N kg ha^−1^ y^−1^) ^[57–59]^. The decline in the soil bacterial diversity due to N-amendment might also facilitate the predominance of nitrophilous species and competitive exclusion of other members as observed (Acidobacteria, Actinobacteria, Firmicutes, etc.). Excessively N-amended paddy soil has been reported to show an increase in nitrophilic species such as Nitrospira ^[60]^, members of Burkholderia and Myxococcales (Anaeromyxobacteraceae) ^[61]^, which was also observed in this study (and discussed below). Zeng et al. 2016 ^[62]^ observed that the N-amendment affected soil microbial diversity through direct nutrient effects and changes in soil and plant properties. Therefore, this study underscored the change in bacterial abundance and diversity among all the treatments, which could be attributed to the influence of the N-amendment.

### Alterations in the bacterial community structure and community response

A notable alteration in the bacterial community structure was observed in this study. Diverse microorganisms thrived in N-amended conditions despite the adverse effects of N. The most abundant bacterial taxa detected in this study, *i.e.,* Gammaproteobacteria, Alphaproteobacteria, Bacilli (Firmicutes), Gemmatimonadota, and Actinobacteria, aligned with previous reports on microbial diversity in As-contaminated paddy soils (Fig. S5 and references herein). Members of these taxa have been previously known for their multifaceted metabolizing ability, which included N₂ fixation, denitrification, and organic matter decomposition. However, the soil microbial community exhibited distinct responses to added N substrates, with some bacterial taxa showing enhanced abundances and others being inhibited, depending on the type of N-amendment. Nevertheless, Sphingomonadales significantly increased across all N-amendments, while Thermosulfovibrionia and Clostridales were negatively impacted. Members of Sphingomonadales have been previously reported for their versatile N metabolizing abilities, improving soil and plant health and facilitating beneficial microbial interactions ^[63–66]^.

In this study, higher N levels (NO_3_^−^ and NO_2_^−^) increased the abundance of Anaerolineae, Dehalococcoidia, and Nitrospirales. Both Anaerolineae and Dehalococcoidia have formerly demonstrated metabolic plasticity in NO_3_^−^ and NO_2_^−^ metabolism, essential for their ecological roles in bioremediation and nutrient cycling in anoxic ecosystems ^[67–71]^. Nitrospirales were favored in NO_3_^−^ and NO_2_^−^ amended sets as observed in this study, which is in line with the previous reports where Nitrospirales significantly impacted N cycling through NO_3_^−^ and NO_2_^−^ metabolism ^[72–74]^. The favorable response of Burkholderia and Gemmatimonadales towards NH_4_^+^ and urea may be linked to their ability of NH_4_^+^ oxidation and urea hydrolysis ^[75, 76]^. Conversely, Pseudomonadales and Kryptoniales thrived at low N concentrations but were negatively affected by higher N levels. Thus, the different responses of the major taxa to the N-amendment seemingly changed the bacterial community structure.

### Impact of different N-amendments on the core microbial community

A core microbiome was identified in all N-amended sets, indicating their crucial role in nitrogen metabolism ^[77]^. The predominant OTUs in this core community were also abundant in the total community, mirroring findings observed in native soil ^[32]^. Core community analysis of the N-amended sets reflected the dominance of Gammaproteobacteria, Alphaproteobacteria, Bacilli (Firmicutes), Gemmatinonadota, Thermoleophilia, Chloroflexi, and Actinobacteriota. Previous studies highlighted the ecological importance of these taxa in nutrient cycling ^[32, 77–80]^. Compared to native soil, a decrease in observed OTUs was evident, with a core community revealing 68% taxonomic variation. Notably, the two most abundant members of the native soil core-community, Nitrosomonadaceae and Beijerinckiaceae diminished, while metabolically versatile Bacillaceae, Pseudomonadaceae, and Sphingomonadaceae emerged as more prominent and resilient under N-amendment. Elevated N resources likely diminished symbiotic interactions among native soil core members, altering canonical N pathways ^[79]^. Nevertheless, the predominance of Bacillaceae, Pseudomonadaceae, Sphingomonadaceae, Gemmatimonadaceae, Paenibacillaceae, Nitrospiraceae, and others in the N-treated community might imply their metabolic versatility in utilizing various C, N, and S sources. Therefore, this study highlights that soil core microbiome can respond dynamically to changes in N-amendment.

### Co-occurrence network of microbial community

The co-occurrence analysis revealed a complex network of microbial community interactions with both positive and negative correlations. It represented strong positive interactions among core community members, including unclassified Alphaproteobacteria, Candidatus Udaeobacter, Azospirillales, Roseiflexaceae, Sphingomonadaceae, and Mycobacterium, which have been reported to be crucial for the nitrogen (N) cycle and microbial dynamics in paddy soil ^[71, 81]^. Alphaproteobacteria, identified as keystone organisms, have been previously known to contribute to N_2_ fixation, enhancing soil fertility and plant growth ^[82]^. The versatile ability of Candidatus Udaeobacter and Azospirillales has also been observed in nitrification and denitrification, vital for sustainable paddy cultivation ^[83, 84]^. Conversely, taxa such as Dadabacteriales, Methyloligellaceae, and Burkholderiales_SC-I-84 were negatively affected by N treatment, which might indicate competition among the microbiota ^[85]^. Ren et al., 2020 ^[86]^ have reported that high N conditions reduced the abundance of Steroidobacteraceae and Burkholderiales_SC-I-84, while Dadabacteriales and Methyloligellaceae have been seen to decrease in diversity due to N fertilization.

The response of the total and the core community of the soil might have implied that N-amendments enhanced copiotroph bacteria (Gammaproteobacteria, Betaproteobacteria, Bacteroidetes) and a few other oligotrophs. However, oligotrophs, such as Verrucomicrobia, Chloroflexota, Actinomycetota, and Alphaproteobacteria, thrived under nutrient-limited conditions, acting as keystone taxa during prolonged fertilization ^[87, 88]^.

## Conclusion

The findings of this study reveal that N-amendments significantly impacted paddy soil bacterial communities, leading to reduced abundance, diversity, and richness of both bacterial and archaeal populations. N-amendment altered soil physicochemical properties and shifted the total as well as core microbial community composition. Notably, core taxa abundant in native soil, such as Nitrosomonadaceae and Beijerinckiaceae, declined under N-treated conditions, while resilient groups like Bacillaceae and Pseudomonadaceae became more prominent. This study indicated a complex interplay among microbial taxa and highlighted the ability of Sphingomomndales to thrive under enhanced nitrogen conditions. Members of Thermodesulfovibrionia and Clostridiales were inhibited under such conditions. Co-occurrence network analysis highlighted the adaptive strategies of soil microbes, where copiotrophic bacteria flourished due to N-amendments, while oligotrophs persisted as keystone taxa even under nutrient-limited conditions. Overall, these insights underscore the delicate balance within paddy soil ecosystems, particularly focusing on the complex structure of the soil microbial community and how N-amendments negatively affect the soil microbial communities. This knowledge can be instrumental in developing sustainable farming methods that not only maintain soil fertility and crop productivity but also preserve the delicate balance of soil microbial life.

## Data accessibility

Raw reads obtained from 16S rRNA gene amplicon submitted in the NCBI (BioProject number PRJNA1000929) database.

## Credit authorship contribution statement

Shushishloka Chakraborty: Conceptualization, Methodology, Software, Validation, Formal analysis, Investigation, Writing – original draft, Writing – review and editing. Pinaki Sar: Conceptualization, Resources, Manuscript Writing, review and editing, Supervision, Project administration, Funding acquisition.

## Acknowledgments

The authors wish to thank Rajendra P. Sahu and Debarshi Mukherjee for their generous help in amplicon sequence analysis techniques, Dr. Himadri Bose and Anindya Sundar Dey for their generous help in sample collection, Dr. Himadri Bose for initiation of microcosm setup, Subhradeep Samaddar, for his assistance in various analyses. Shushishloka Chakraborty is thankful for the financial support from the Ministry of Human Resource Development, Government of India, as her doctoral fellowship.

## SUPPORTING MATERIALS

**Figure 1S.**
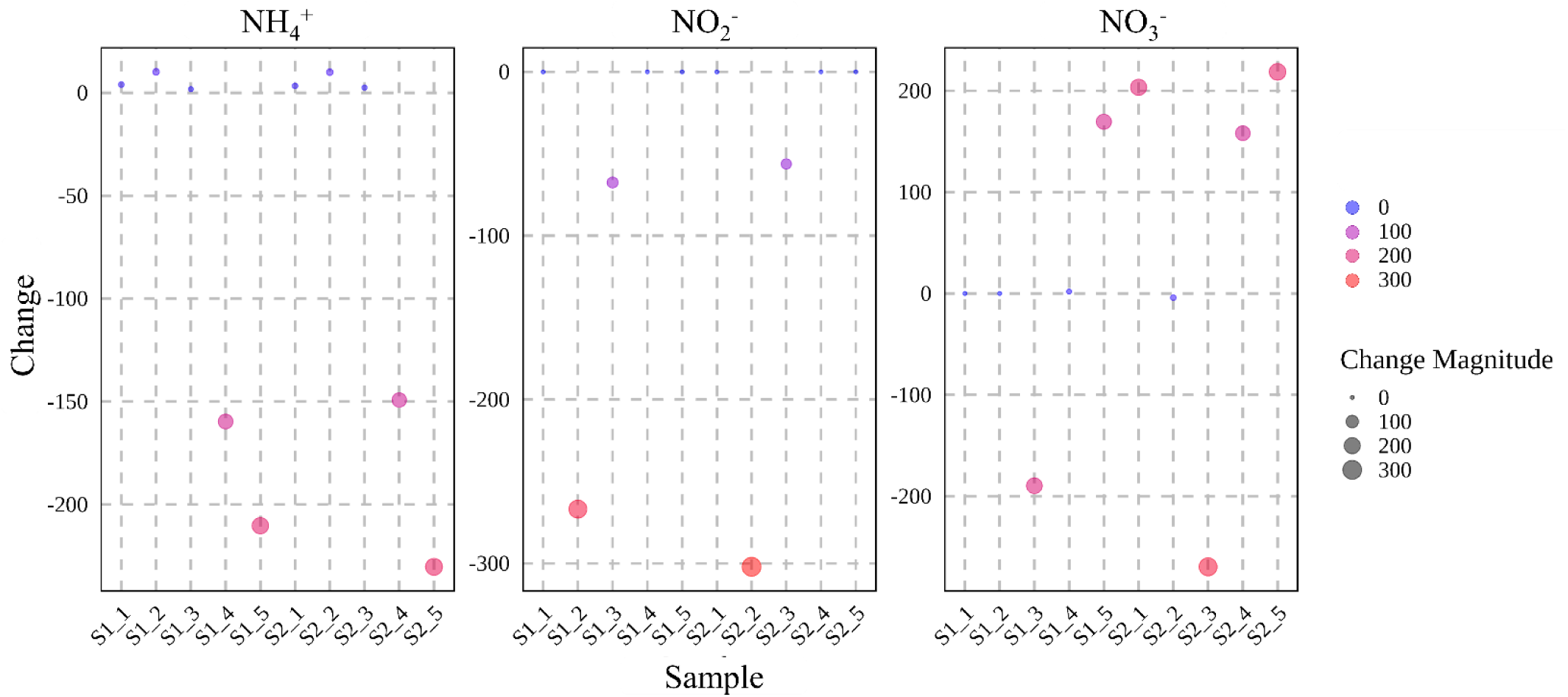
Change in concentration of major N sources before and after incubation

**Figure 2S.**
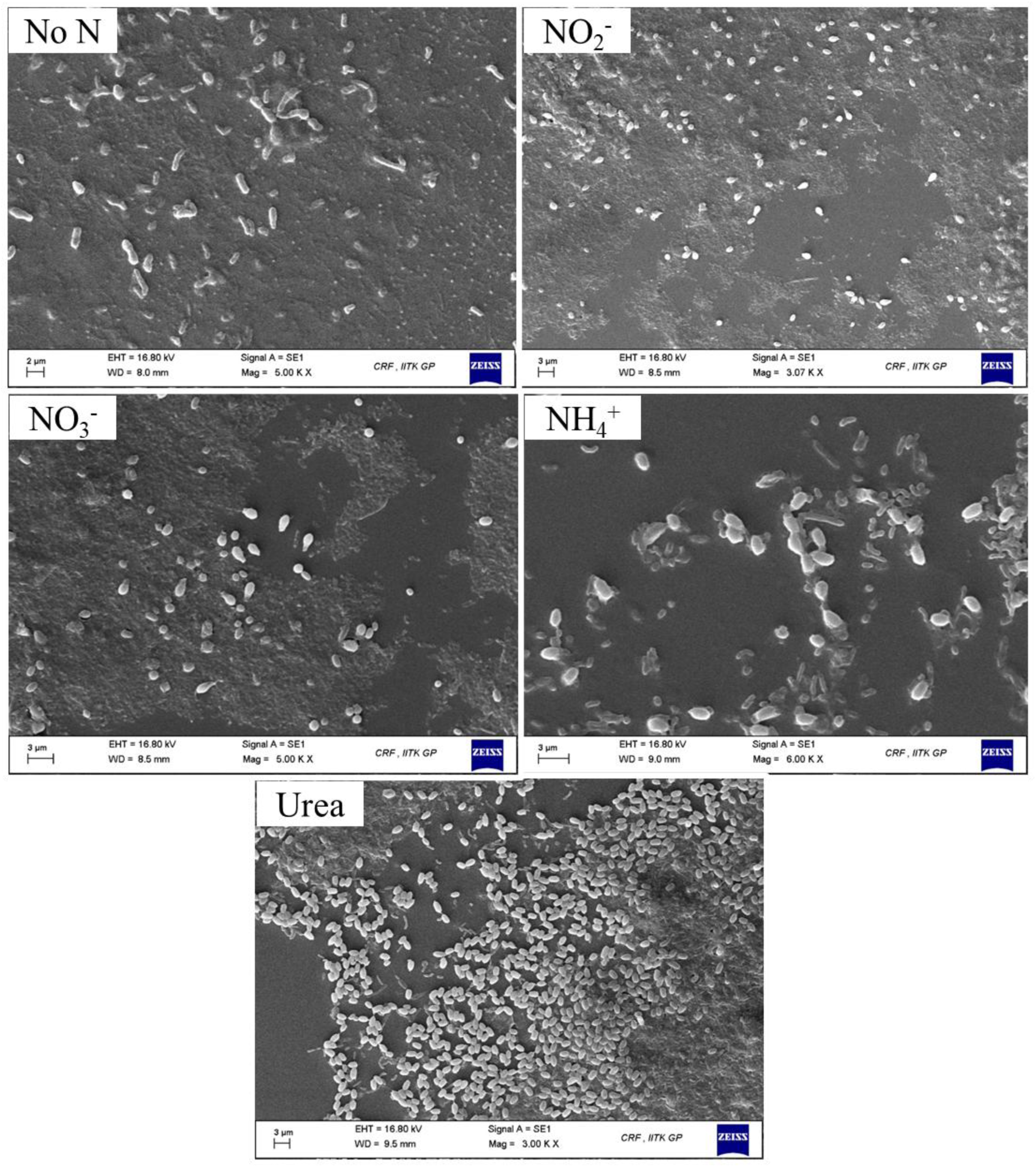
Scanning electron microscopic images of microbial cells from N un-treated and -treated communities [No N added, NO_2_^−^, NO_3_^−^, NH_4_^+^, Urea]

**Figure 3S.**
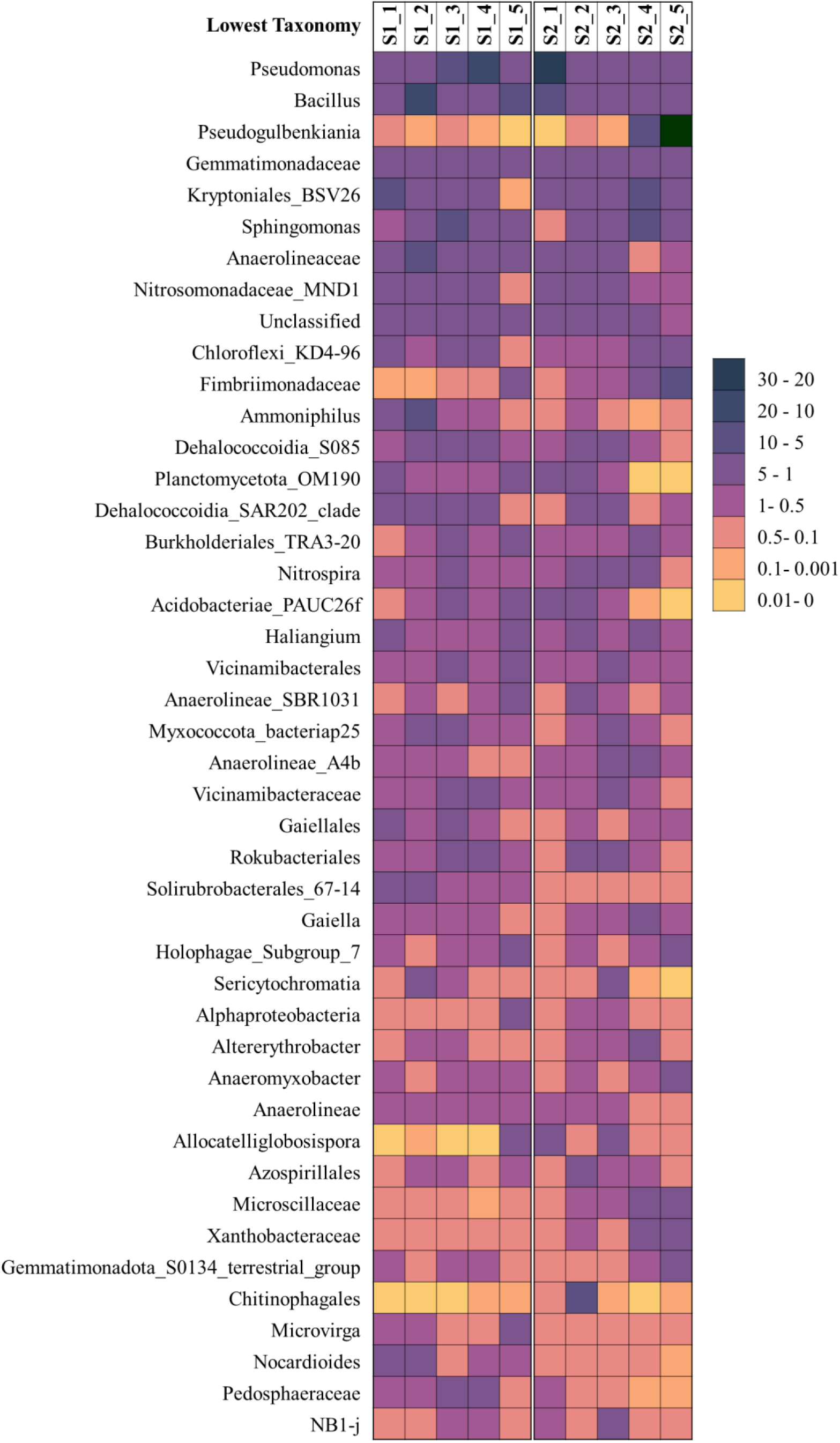
Heatmap displaying relative abundance of major taxa (at lowest taxa level, average relative abundance > 0.1 %) of the total community members

**Figure 4S.**
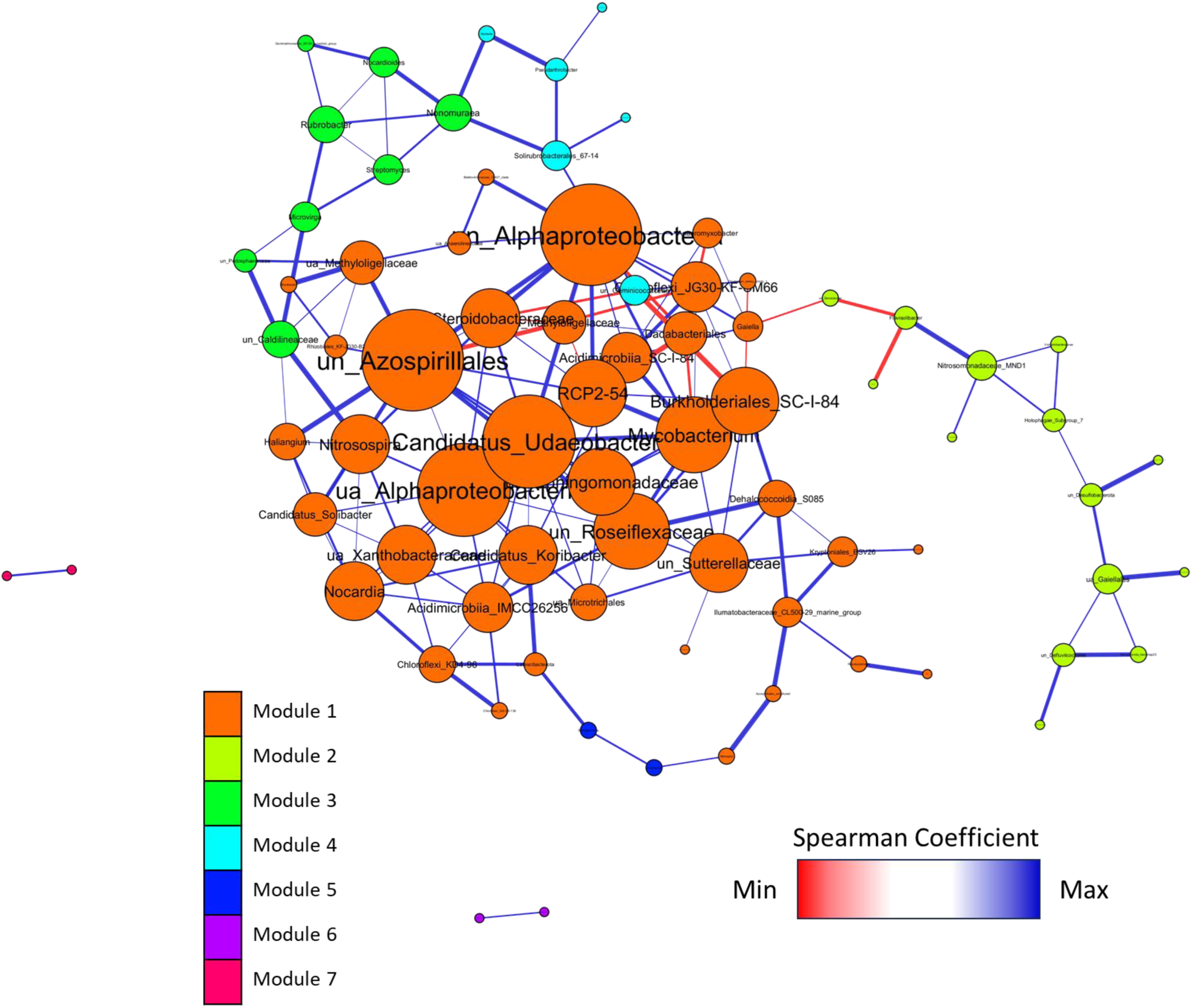
Co-occurrence network of the core microbial community (lowest taxonomy level). Node size is proportional to the number of degrees; nodes with higher degrees are bigger in size than those with lower connections. The color of the nodes indicates the modules classifying the taxa in & different modules. The color of the edges represents the strength of Spearman’s correlations between the nodes. The strength of the connections is indicated in blue as being stronger and red as being weaker.

**Figure 5S.**
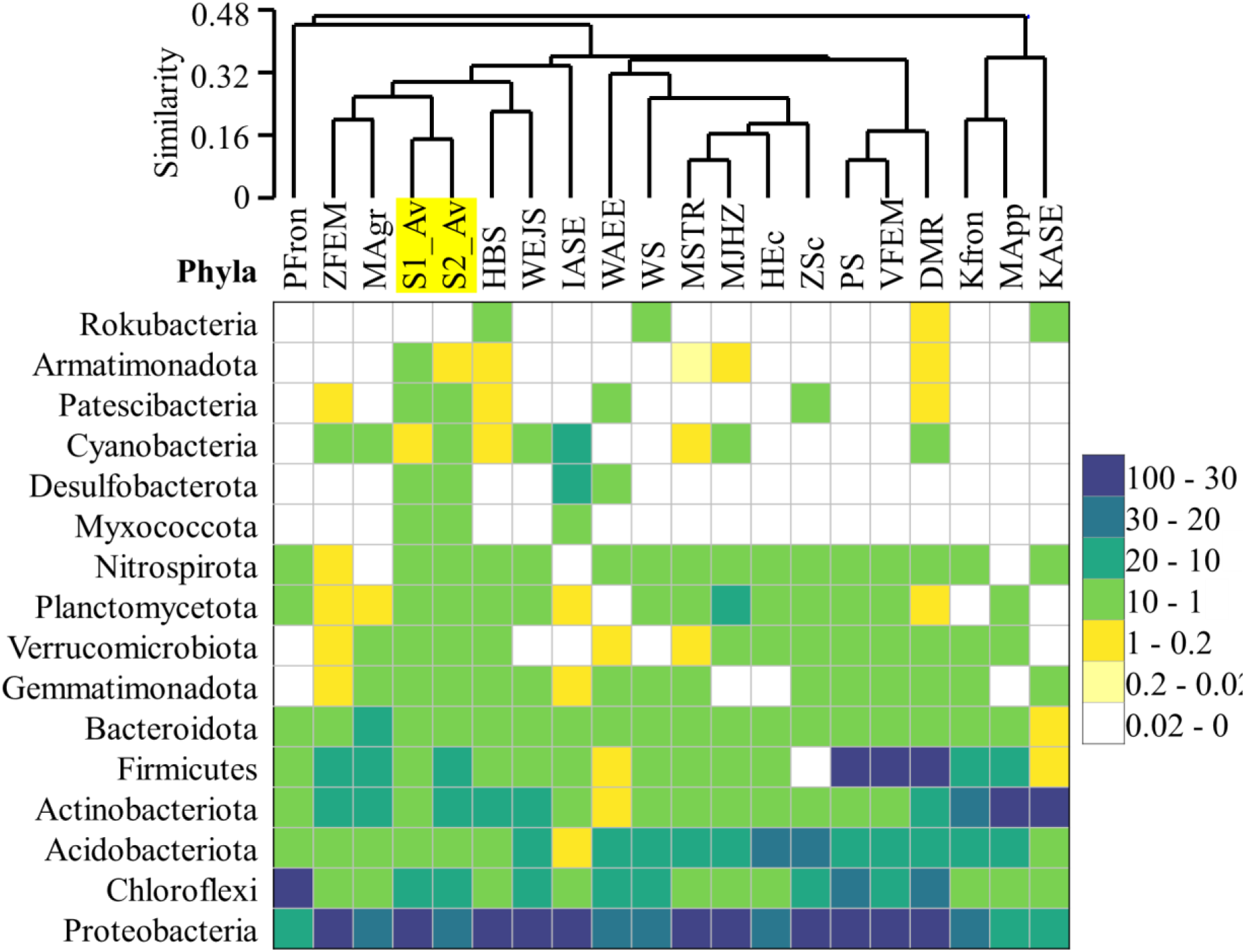
Heatmap and hierarchical clusters on bacterial communities, as well as various other paddy soils, were detected in this study. The geological location and details of each sample are presented in a table (Table S5). Details of the taxonomic data are provided in Supplementary File 1

**Table S1.** Concentration of major N sources before and after incubation.

| Sample ID | NO<sub>2</sub><sup>-</sup> 0day | NO<sub>2</sub><sup>-</sup> final | NO<sub>3</sub><sup>-</sup> 0day | NO<sub>3</sub><sup>-</sup> final | NH<sub>4</sub><sup>+</sup> 0day | NH<sub>4</sub><sup>+</sup> final |
| --- | --- | --- | --- | --- | --- | --- |
| S1 1 | 0.0 | 0.0 | 0.0 | 0.0 | 0.8 | 4.8 |
| S1 2 | 266.9 | 0.0 | 0.0 | 0.0 | 0.1 | 10.3 |
| S1 3 | 67.6 | 0.0 | 212.8 | 23.1 | 0.3 | 2.1 |
| S1 4 | 0.0 | 0.0 | 0.0 | 1.8 | 160.8 | 1.0 |
| S1 5 | 0.0 | 0.0 | 0.0 | 169.5 | 211.6 | 1.3 |
| S2 1 | 0.0 | 0.0 | 0.0 | 203.5 | 0.3 | 3.7 |
| S2 2 | 302.1 | 0.0 | 4.9 | 0.8 | 0.8 | 10.9 |
| S2 3 | 56.2 | 0.0 | 280.7 | 11.0 | 0.2 | 2.6 |
| S2 4 | 0.0 | 0.0 | 0.0 | 158.1 | 150.2 | 0.9 |
| S2 5 | 0.0 | 0.0 | 0.0 | 218.7 | 231.2 | 0.9 |

**Table S2.** Relative abundance of major taxa (Phylum – Class levels) of the total microbial communities, phyla having average abundance < 1% in the total community were grouped into ‘Others’ [This data is used to plot Fig 2a].

| Phylum-Class | S1 1 | S1 2 | S1 3 | S1 4 | S1 5 | S2 1 | S2 2 | S2 3 | S2 4 | S2 5 |
| --- | --- | --- | --- | --- | --- | --- | --- | --- | --- | --- |
| Gammaproteobacteria | 9.62 | 8.94 | 20.61 | 26.91 | 12.46 | 29.49 | 15.68 | 16.91 | 19.30 | 41.39 |
| Alphaproteobacteria | 9.10 | 10.74 | 15.34 | 9.21 | 16.59 | 7.75 | 12.83 | 9.84 | 19.25 | 10.51 |
| Bacilli | 7.41 | 20.31 | 4.54 | 2.22 | 9.88 | 9.04 | 4.16 | 5.25 | 4.05 | 1.77 |
| Anaerolineae | 5.14 | 9.78 | 4.73 | 4.15 | 7.92 | 5.19 | 8.50 | 7.86 | 3.07 | 3.84 |
| Bacteroidota | 10.33 | 2.90 | 3.23 | 6.20 | 2.16 | 2.59 | 9.93 | 6.38 | 10.08 | 5.89 |
| Acidobacteriota | 4.73 | 4.40 | 6.93 | 5.03 | 8.17 | 5.64 | 7.99 | 7.89 | 4.38 | 4.28 |
| Gemmatimonadota | 4.51 | 3.53 | 5.58 | 5.86 | 4.30 | 3.01 | 4.55 | 4.49 | 5.24 | 5.24 |
| Myxococcota | 5.67 | 4.42 | 5.68 | 4.43 | 4.43 | 3.92 | 5.69 | 4.68 | 4.25 | 2.95 |
| Actinobacteria | 5.79 | 6.95 | 2.59 | 3.83 | 6.84 | 3.31 | 2.69 | 2.40 | 2.81 | 2.50 |
| Thermoleophilia | 6.37 | 4.07 | 3.52 | 3.90 | 2.08 | 1.52 | 2.50 | 1.67 | 3.16 | 2.83 |
| Verrucomicrobiota | 1.78 | 1.77 | 1.95 | 2.67 | 4.61 | 1.68 | 2.94 | 2.69 | 2.73 | 1.47 |
| Dehalococcoidia | 2.00 | 2.12 | 4.28 | 2.18 | 1.01 | 1.01 | 3.15 | 3.02 | 1.21 | 1.04 |
| Desulfobacterota | 1.88 | 0.64 | 0.69 | 2.71 | 1.41 | 6.15 | 0.48 | 2.66 | 3.01 | 1.04 |
| Planctomycetota | 2.77 | 1.29 | 1.51 | 1.61 | 3.38 | 2.19 | 2.70 | 2.79 | 0.12 | 0.17 |
| Patescibacteria | 1.54 | 1.36 | 2.96 | 3.65 | 0.95 | 1.76 | 2.10 | 1.76 | 1.05 | 0.47 |
| Armatimonadota | 0.54 | 0.63 | 1.19 | 0.33 | 2.23 | 0.63 | 1.22 | 1.19 | 3.62 | 5.94 |
| Unassigned | 1.48 | 1.28 | 1.64 | 1.67 | 1.53 | 2.41 | 1.92 | 3.41 | 1.07 | 0.68 |
| Clostridia | 2.74 | 1.45 | 0.51 | 1.10 | 2.06 | 2.61 | 0.28 | 1.00 | 0.50 | 0.31 |
| Chloroflexi_KD4-96 | 1.20 | 0.98 | 1.20 | 1.56 | 0.37 | 1.00 | 0.75 | 0.54 | 3.29 | 1.58 |
| Nitrospirota | 1.28 | 0.87 | 1.60 | 0.64 | 0.98 | 1.45 | 1.51 | 1.93 | 1.10 | 0.49 |
| Cyanobacteria | 3.58 | 3.17 | 1.42 | 0.47 | 0.33 | 0.25 | 0.28 | 1.68 | 0.04 | 0.08 |
| Archaea unassigned | 0.06 | 0.06 | 0.06 | 0.03 | 0.07 | 0.06 | 0.06 | 0.05 | 0.05 | 0.12 |
| Aenigmarchaeota | 0.01 | 0.00 | 0.03 | 0.00 | 0.01 | 0.01 | 0.01 | 0.01 | 0.04 | 0.04 |
| Micrarchaeota | 0.00 | 0.00 | 0.00 | 0.00 | 0.00 | 0.00 | 0.00 | 0.00 | 0.00 | 0.00 |
| Thermoplasmata | 0.00 | 0.00 | 0.00 | 0.00 | 0.00 | 0.00 | 0.00 | 0.00 | 0.00 | 0.00 |
| Crenarchaeota | 0.00 | 0.00 | 0.00 | 0.00 | 0.00 | 0.00 | 0.00 | 0.00 | 0.00 | 0.00 |
| Others | 10.47 | 8.32 | 8.23 | 9.65 | 6.24 | 7.36 | 8.08 | 9.90 | 6.59 | 5.36 |

**Table S3.**
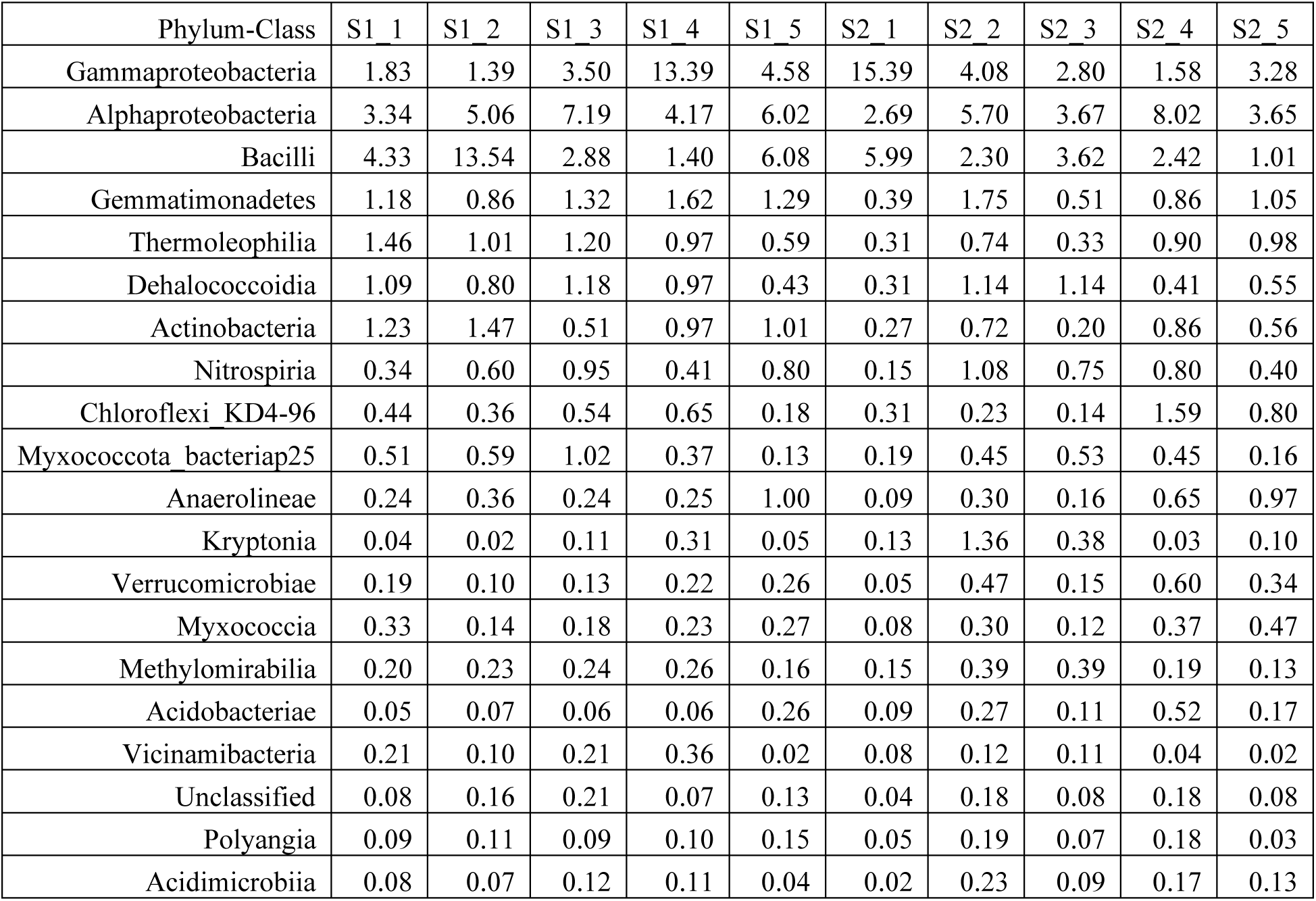
Relative abundance of major taxa (Phylum – Class levels) of the core microbial communities, phyla having average abundance < 0.1% in the total community were grouped into ‘Others’ [this data is used to plot Fig 3b].

**Table S4.** Network topology.

|  | Positive network | Negative network | Module network |
| --- | --- | --- | --- |
| Clustering coefficient | 0.361 | 0 | 0.363 |
| Connected components | 2 | 3 | 3 |
| Network diameter | 13 | 7 | 15 |
| Network radius | 7 | 4 | 8 |
| Network centralization | 0.237 | 0.309 | 0.12 |
| Characteristic path length | 4.997 | 2.909 | 5.176 |
| Avg. number of neighbors | 4.577 | 2.167 | 4.575 |
| Number of nodes | 71 | 17 | 77 |
| Number of edges | 153 | 16 | 169 |
| Network density | 0.064 | 0.197 | 0.064 |
| Network heterogeneity | 0.759 | 0.62 | 0.683 |
| Isolated nodes | 0 | 0 | 0 |
| Number of Self-loop | 0 | 0 | 0 |
| Multi-edge node pairs | 0 | 0 | 0 |
| Analysis time (sec) | 0.01 | 0.04 | 0.05 |

**Table S5.** The geological location of each sample.

| Sample ID | Location | References |
| --- | --- | --- |
| PFron | Dingdian village, Natong Town, China | [71] |
| ZFEM | Castello d'Agogna, Pavia, Italy | [89] |
| S1_Av | Bethuadahari, West Bengal, India |  |
| S2_Av | Bethuadahari, West Bengal, India |  |
| HBS | Bethuadahari, West Bengal, India | [32] |
| WAEE | Jiangsu Province, China | [90] |
| Kfron | Bihar, India | [91] |
| KASE | Tamil Nadu, India | [92] |
| WS | Zhanggong, Jinxian County, China | [93] |
| PS | Sundarban, West Bengal, India | [94] |
| DMR | Dudhali village, Dehradun and Shilimb village, Pune, India | [95] |
| MSTR | Katwa, West Bengal India | [96] |
| MJHZ | Chakdaha block, West Bengal, India | [97] |
| WEJS | Yellow River, Rencheng, China | [98] |
| IASE | Doñana wetlands, Southwest Spain | [99] |
| HEc | Vercelli, Italy | [100] |
| ZSc | Dingshu, China | [101] |
| MApp | Treintay Tres, Uruguay | [102] |
| MAgr | Gongzhuling, Jilin Province, China | [103] |
| VFEM | Vercelli, Italy | [104] |

